# Within-subject changes in methylome profile identify individual signatures of early-life adversity, with a potential to predict neuropsychiatric outcome

**DOI:** 10.1101/2023.12.16.571594

**Authors:** Annabel K. Short, Ryan Weber, Noriko Kamei, Christina Wilcox Thai, Hina Arora, Ali Mortazavi, Hal S. Stern, Laura Glynn, Tallie Z. Baram

**Author notes:** **Corresponding author**: Tallie Z. Baram. **Email:**.

## Abstract

**Background:** Adverse early-life experiences (ELA), including poverty, trauma and neglect, affect a majority of the world’s children. Whereas the impact of ELA on cognitive and emotional health throughout the lifespan is well-established, it is not clear how distinct types of ELA influence child development, and there are no tools to predict for an individual child their vulnerability or resilience to the consequences of ELAs. Epigenetic markers including DNA-methylation profiles of peripheral cells may encode ELA and provide a predictive outcome marker. However, the rapid dynamic changes in DNA methylation in childhood and the inter-individual variance of the human genome pose barriers to identifying profiles predicting outcomes of ELA exposure. Here, we examined the relation of several dimensions of ELA to changes of DNA methylation, using a longitudinal within-subject design and a high threshold for methylation changes in the hope of mitigating the above challenges.

**Methods:** We analyzed DNA methylation in buccal swab samples collected twice for each of 110 infants: neonatally and at 12 months. We identified CpGs differentially methylated across time, calculated methylation changes for each child, and determined whether several indicators of ELA associated with changes of DNA methylation for individual infants. We then correlated select dimensions of ELA with methylation changes as well as with measures of executive function at age 5 years. We examined for sex differences, and derived a sex-dependent ‘impact score’ based on sites that most contributed to the methylation changes.

**Findings:** Setting a high threshold for methylation changes, we discovered that changes in methylation between two samples of an individual child reflected age-related trends towards augmented methylation, and also correlated with executive function years later. Among the tested factors and ELA dimensions, including income to needs ratios, maternal sensitivity, body mass index and sex, unpredictability of parental and household signals was the strongest predictor of executive function. In girls, an interaction was observed between a measure of high early-life unpredictability and methylation changes, in presaging executive function.

**Interpretation:** These findings establish longitudinal, within-subject changes in methylation profiles as a signature of some types of ELA in an individual child. Notably, such changes are detectable beyond the age-associated DNA methylation dynamics. Future studies are required to determine if the methylation profile changes identified here provide a predictive marker of vulnerabilities to poorer cognitive and emotional outcomes.

**Funding:** Supported by NIH P50 MH096889, a Precision Medicine Initiative grant from the State of California (OPR20141) and the Bren Foundation.

**Research in context:** *Evidence before this study:* Identification of individuals at risk for cognitive and emotional problems is required for targeted interventions. At the population level, experiencing early-life adversity has been consistently linked to an elevated susceptibility to various mental illnesses. However, recent studies have revealed a significant limitation in the ability of early-life adversity to predict individual-level risk, and there is presently no reliable tool available to determine whether a child experiencing adversity will develop future mental health problems. Promising efforts to discover predictive markers by examining DNA methylation in peripheral cells are challenged by extensive genetic and epigenetic population variability and the rapid methylation changes taking place during childhood, rendering the identification of clinically valuable predictive markers a complex endeavor.

*Added value of this study:* This study examined neurodevelopmental outcomes following several dimensions of ELA, including a recently identified dimension-unpredictability of parental and environmental signals to the child. It demonstrates changes in DNA methylation in children exposed to a spectrum of ELA dimensions and severity using alternative approaches to those used previously: It employs a longitudinal within-subject design, enabling assessment of DNA changes within an individual over time rather than a cross section comparison of different groups, and focuses on the first year of life, an understudied epoch of development. The study uses reduced representation bisulfite sequencing to measure methylation, an approach compromising between targeted sequencing and a whole genome approach, and sets a high threshold for methylation changes, in consideration of the large changes of DNA methylation during childhood. Finally, in accord with emerging discoveries of the differential effects of ELA on males and females, the study uncovers sex-effects arising already before puberty.

*Implications of all the available evidence:* Collectively, our study, together with a robust existing literature (1) identifies early-life unpredictability as an additional determinant of DNA methylation changes, (2) indicates that within-subject changes in methylation profiles of peripheral cells hold promise as precision medicine tools for predicting risk and resilience to the adverse consequences of early-life hardships on mental health, and (3) suggests that sex-differences should be explored even prior to puberty. Our study contributes significantly to the important goal of early identification of predictive “epigenetic scars” caused by adverse early-life experiences. Such markers are required for targeting interventions to those most at need.

## Introduction

Early-life experiences may exert a profound cumulative impact on lifespan trajectories of mental and physical health. Unsurprisingly, a robust body of work has focused on the contribution of salient early-life experiences, and especially of early-life adversity (ELA) to cognitive and mental health outcomes. In cohorts from diverse countries, socioeconomic levels and cultures, such studies have typically focused on the level of adversity, the cumulative impact of different types of adversity, and the distinct impacts of dimensions of adversity such as deprivation vs threat (1–14). This strong literature supports the roles of ELA and its specific dimensions in predisposing individuals to physical, cognitive and mental health disorders (15–17). However, the predictive value of ELA to vulnerability to mental and physical health problems applies well at the population level, whereas ELA exposures predict the outcome of an individual child little better than chance (18). Therefore, a significant unmet challenge remains in our ability to predict for an individual child whether they will be vulnerable or resilient to physical, cognitive or mental health problems.

Whereas trauma, poverty and abuse early in life significantly increase the risk of experiencing poorer cognition and mental health throughout the lifespan (19–25), the significant amount of variance unaccounted for in child developmental outcomes has led to a search for additional potential sources of adversity that might have been missed. An additional dimension of adversity, which explains some of the variance in cognitive and emotional outcomes, is unpredictability of the early parental care behaviors and home environment. Initially detected in experimental animal models of early-life adversity (26–28), unpredictable sequences of parental care behaviors have emerged as an important potential predictor of susceptibility to later cognitive and emotional deficits (29). Specifically, in experimental rodent models, resource scarcity elicited fragmented and unpredictable sequences of maternal care during a sensitive developmental period. In turn, these aberrant sensory signals to the developing brain influenced brain circuit maturation by altering selective microglial pruning of neuronal synapses (30), leading to significant impairments of cognitive functions (31–33) and reward behaviors (27,34–36). In human studies, this additional novel ELA was characterized initially by measuring unpredictable sequences of sensory signals that the caregiver transmits to the infant and child (such as speech, touch or visual cue). The concept was then extended to other proximate sources of signals to the developing brain, including, in addition to caregivers, also the household and environment. The multiple sources and timescales of unpredictability are measured using the Questionnaire on the Unpredictability of Childhood (QUIC) (37,38). The contribution of unpredictable signals from caregivers and home environment (“unpredictability”) to children’s cognitive and emotional functions has now been established across diverse populations (37,39–44), and remains robust upon inclusion in the statistical models of well-established adverse experiences. Unpredictable parental and environmental signals to the infant impact the maturation of structural and functional brain connectivity, measured using magnetic resonance imaging (45,46), and predict emotional problems also in adulthood (47).

Sex differences in the outcomes of ELA have been recognized, with women who endorse depression, anxiety or addiction being more likely to report adverse early-life experiences compared with men (48–58). However, whether or not such sex differences emerge prior to puberty and whether females are more vulnerable already in childhood has not been resolved. Our studies on the consequences of early-life unpredictability are now uncovering selective vulnerability in girls (59), providing an impetus to examine the role of sex in the current studies.

Studies focusing on predicting the impact of ELA on later life mental and physical health involve three elements: the nature of the insult(s), an appropriate, universally applicable outcome, and an accessible reliable marker, detectable already early in life, that correlates robustly with the outcome measures. Here we looked at several dimensions of ELA, including unpredictability, as ‘drivers’ of neurodevelopmental outcomes. We chose children’s ability to regulate behavior and attention as an outcome measure, because it is associated with having a high level of executive function (15,60–63). In turn, executive function during childhood is one of the most robust predictors of cognitive and emotional outcomes and of success throughout life. A child’s self-regulation abilities (effortful control) emerge towards the end of the first year of life and continue to develop throughout childhood. Importantly, effortful control is highly influenced by early-life experiences (36, 47). Therefore, in the current study we focused on effortful control as a key neuro-developmental outcome.

We employed changes in DNA methylation as a potential marker of (‘signature’) ELA, including unpredictability, because such changes are known to be highly sensitive to environmental influences (65,66). The levels of methylation of specific DNA nucleotides and their changes throughout development have been a topic of extensive study (67–75). Indeed, DNA methylation patterns correlate with chronological age, providing ‘epigenetic’ or ‘DNA-methylation’ clocks (67,68,72). Adversity throughout life (74,76–79), as well as mental and physical disease states (80) have been shown to accelerate this ‘epigenetic age’, and even to predict the timing of death (69,77).

However, the use of DNA methylation as a ‘signature’ of ELA and a potential predictor of health and development has been challenging. Whereas at the population level, DNA methylation ‘signatures’ of adversity / stress are apparent at both individual timepoints and across time (73,74,79), their role as a predictive marker for an individual child has been limited by the high levels of inter-individual differences of the human genome and its DNA methylation patterns. In addition, the development of methylation risk scores for the outcomes of ELA needs to consider the highly dynamic changes of DNA methylation early in life: Changes in DNA methylation levels during childhood are around 4 fold more rapid than in adults (81) and involve both augmented methylation as well as demethylation, depending on which CpG sites are examined (75,76,81). Thus, identifying DNA methylation patterns and changes that associate with ELA and predict outcome for a given child has continued to present a challenge, potentially requiring setting robust criteria for methylation changes, which allow detection of adversity effects beyond those of age (72,73,76,79,81).

Here we capitalized on our preclinical studies that had employed a within-subject design and a high-threshold criterion for DNA methylation changes. Those studies allowed distinguishing the effects of ELA from those of age (82). In the current study we examined DNA methylation changes between two samples obtained from the same child (in the neonatal period and at one year of age) and assessed methylation changes with a high change threshold. We then correlated these changes with several dimensions of ELA as well as with the outcome measure of executive function at age 5 years. We identify a novel set of DNA methylation changes that correlate with outcomes as well as with unpredictability in a sex-specific manner and may thus be useful as a predictive marker in future studies.

## Methods

### Participants

Study participants were 126 infants enrolled before birth, part of a longitudinal study evaluating the role of early experiences in cognitive and emotional development. All study procedures were approved by the Institutional Review Board for Protection of Human Subjects at Chapman University and the University of California-Irvine. Each mother provided written, informed consent for herself and her child. Demographic information for the cohort appears in Table 1.

**Table 1.**
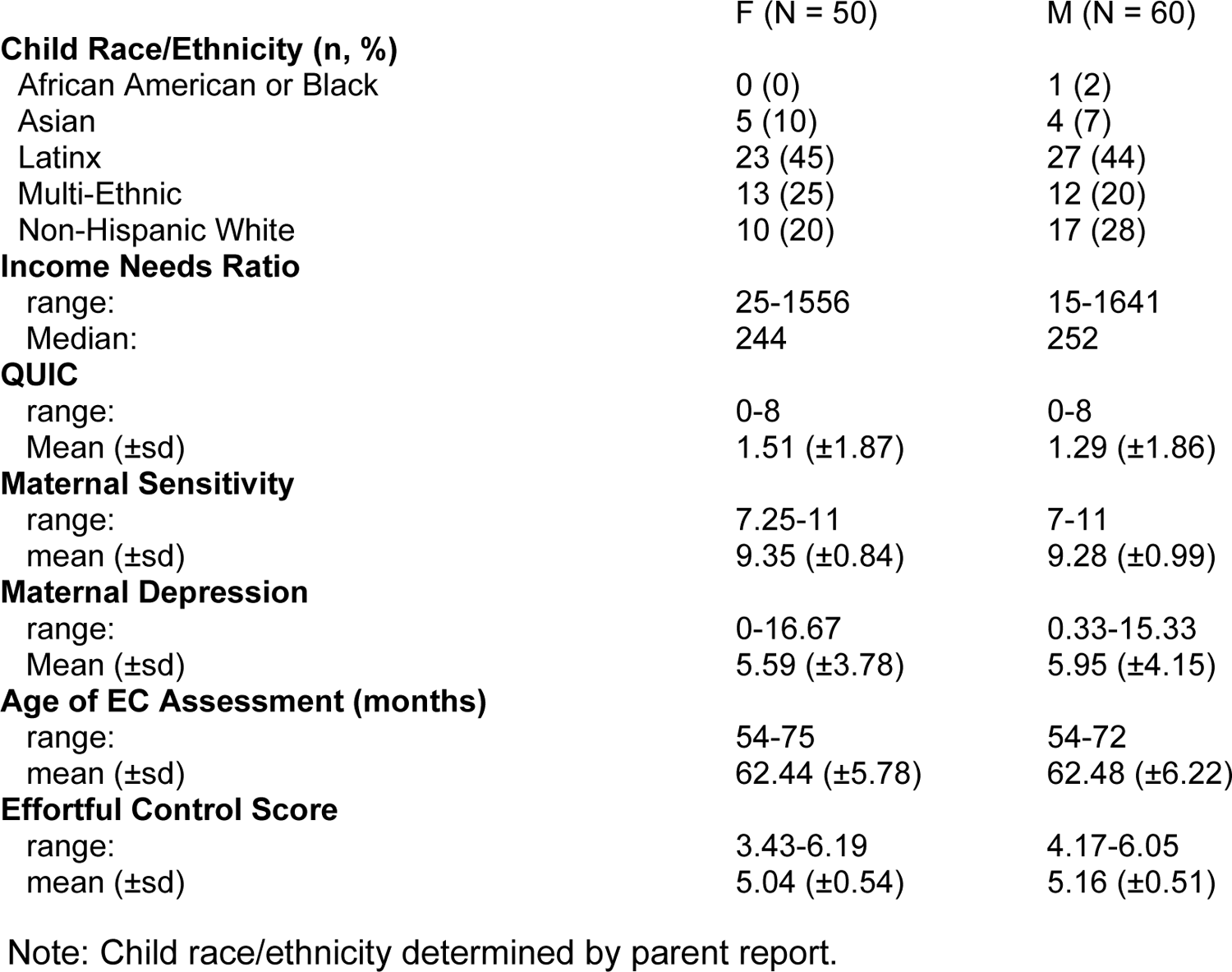
Demographic characteristics of sample.

Five samples were removed because of low sequencing reads (<9 million), seven samples were excluded due to high variability in the number of sequenced sites, and four samples were removed after participants were diagnosed with significant learning impairments. Analysis was performed on remaining 110 samples (m=60, f=50)

### Income to Needs Ratio

Family income-to-needs ratio was calculated by dividing total annual household income by the appropriate U.S. Census Bureau poverty threshold based on family size. The median income-to-needs ratio in this sample was 2.44 (244%), and according to federal guidelines, those families living below 200% of the federal poverty line (a ratio of 2.0) are considered low income and would, for example, qualify for the Supplemental Nutrition Assistance Program (SNAP). However, the families included in our cohort reside in Southern California, which is a relatively expensive area of residence, while the income-to-needs ratio standards are based on national living standards. Therefore, using metrics that adjust for the cost of living in the county of residence (83), a median income to needs ratio of 2.44 corresponds to 70% of the families living below the living wage level.

### Maternal Sensitivity

At 6 and 12-months postpartum, maternal sensitivity was evaluated using a coding scheme developed for the National Institute for Child Health and Development (NICHD) Study for Early Child Care and Youth Development (84). This paradigm is an objective, behaviorally-based laboratory assessment tool for studying maternal behavior that is well-validated and predictive of the quality of mother-child attachment (84,85). Mother-child pairs were videotaped in a semi-structured 10-minute play session, in which mothers are given a standard set of age-appropriate toys and told to play with their infant as they would at home. Following the NICHD procedure (86), a composite rating of quality of maternal care is created by summing ratings of sensitivity to non-distress, maternal positive regard, and intrusiveness (reverse-coded). Twenty percent of the tapes were selected at random, without coder knowledge, and double-coded to obtain an index of inter-rater reliability, which averaged 88% across the two assessments. The correlation between 6 and 12 months was 0.36 and so they were combined to create a single composite measure, to provide a consistent measure of this dimension of maternal care throughout the first postnatal year. Separate analyses of the correlation of maternal sensitivity at 6 months or 12 months with effortful control are provided in the Supplemental Materials (Figure S-3).

### Maternal Depressive Symptoms

At 2, 6, and 12 months postpartum maternal depressive symptoms were measured with the 10-item Edinburgh Postnatal Depression Scale (EPDS) (87). Possible scores on this scale range from 0 to 30, with a score of 10 or more indicating probable minor depression and 13 or more likely major depression. In the present study, 38 percent of the mothers scored above the threshold for minor depressive episode and 18 percent scored in the probable major depressive range for at least one of the postpartum assessments. Correlations between the depression scores at the three timepoints ranged from 0.57 to 0.66. In view of these high correlations, the scores at the three time-points for each mother were averaged to provide a consistent measure of maternal depressive symptoms throughout the first postnatal year.

### Unpredictability

Unpredictability of signals from the caretaker(s) has been identified by us as an important dimension of early life adversity (37–41), and these findings have been confirmed and extended by others (42). Here, we assessed unpredictability of the early environment with the Questionnaire of Unpredictability in Childhood (QUIC;(37)). The original self-report version of the QUIC is a 38-item questionnaire that assesses exposure to unpredictability in social, emotional and physical domains of a child’s environment. It displays excellent psychometric properties (α = 0.89; test-retest reliability = 0.91) and is associated with observational measures of parental and household unpredictability. For the purposes of this study, the parent-report preschool version of the QUIC was employed (88). This version predicts both behavioral observations and parent-report of child effortful control (41,59). Items and endorsement rates are included as Supplemental Table S-1.

### Child Effortful Control

Effortful control is widely considered an optimal measure of executive function in children. Notably, in large prospective studies, effortful control in young children was an excellent predictor of school performance and of success later in life (15,60–63). Here, at roughly five years of age, effortful control measured with the Child Behavior Questionnaire (CBQ) (89) was collected from 90 participants who attended the 5 year follow up visit. This measure exhibits strong internal reliability and validity (90–92) and consistency between parent report and home and laboratory observations (93,94).

### Buccal swab collection

Infant DNA samples were collected via buccal swab from newborns (M_age_ = 2.6 weeks, SD = 0.92) and again at one year (M_age_ = 12.4 months, SD = 0.52) using the DNA Genotek Oragene Discover (DNA Genotek Cat# OGR-575) kit.

### Isolation and quantification of DNA for making reduced representation bisulfite sequencing (RRBS) libraries from Human Buccal swab/saliva

The Buccal swab/saliva samples were incubated at 50°C for 2 hours. Next, 1/25 volume of prepIT-L2P (DNA Genotek Cat# PT-L2P-45) was added, samples were incubated on ice for 10 min and centrifuged at room temperature to collect the supernatant. Genomic DNA was prepared from this supernatant using the Quick gDNA kit (Zymo Research, Cat# D3025) following the manufacturer’s protocol. The quantity of double-stranded DNA was analyzed using Qubit.

RRBS libraries were prepared from 200 ng of genomic DNA digested with MspI restriction enzyme and then extracted with ZR-DNA Clean & Concentrator™-5 kit (Zymo Research, Cat# D4014). According to Illumina’s specified guidelines, fragments were ligated to pre-annealed adapters containing 5’-methylcytosine instead of cytosine (www.illumina.com). Adaptor-ligated fragments were then bisulfite-treated using the EZ DNA Methylation-Lightning™ Kit (Zymo Research, Cat# D5459). Preparative-scale PCR (16 cycles) was performed with Illumina index primers, and the resulting products were purified with DNA Clean & Concentrator for sequencing. Amplified RRBS libraries were quantified and qualified by Qubit, Bioanalyzer (Agilent), and Kapa library quant (Kapa systems, Cat# 07960140001) and then sequenced with paired-end 100 bp on the Illumina Nova-seq platform. Based on initial experiments, we chose a depth of 25 million for newborns and 50 million for one-year-olds to gain an average of 10 million mapped reads on the human genome for all samples. Samples were sequenced in batches and no batch control was employed.

### RRBS processing and detection of differentially methylate sites (DMSs)

The overall workflow and analytical pipeline is depicted in the schematic below:

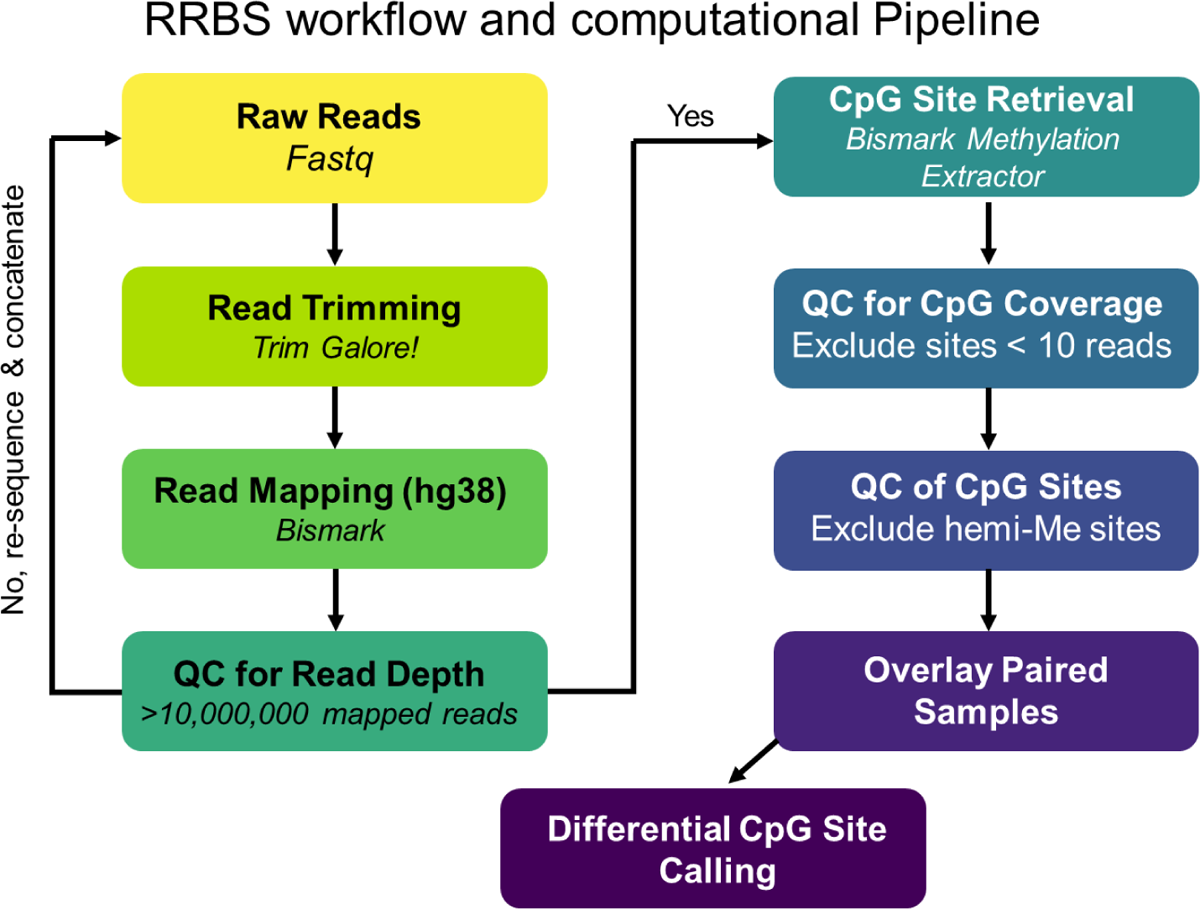

Adaptor and low-quality reads were trimmed and filtered using Trim Galore! 0.4.3 (http://www.bioinformatics.babraham.ac.uk/projects/trim_galore/, RRID:SCR_011847) with the parameter “--fastqc –stringency 5–rrbs –length 30 –non_directional.” Reads were aligned to the human genome (hg38) by using Bismark 0.16.3 ((95) RRID:SCR_005604) with “---non_directional” mode. CpG sites were called by “bismark_methylation_extractor” function from Bismark. Sites with coverage of more than 10 were accepted for further study. Differential methylation sites (DMSs) were first called using Methylkit (R 4.0.5) ((96) RRID:SCR_005177) to identify sites with a minimum ±5% change (82) between sample A (neonatal) and B (one year of age) using single CpG sites. We chose a 5% methylation change as such a change distinguished the influence of ELA in a preclinical study (82), and because it is larger than is expected from changes related to age alone during the first year of life, when DNA methylation is rapidly changing (75,76,81). A site that was identified as having a minimum 5% change in methylation in any individual was then significance-tested for a change in methylation between the A and B samples resulting in a p-value for the i-th site of pi. A test was then carried out to determine sites with a significant change from the A to B samples across all individuals by combining the individual test results using −2 ∑^n^_i=1_ ln (*p_i_*) as a test statistic as described in (97). A site that passed a Benjamini-Hochberg false discovery rate of q=0.1 with a cut off = 0.00005 was determined to be a DMS.

### Calculation of DNA methylation level/percentage and delta methylation

The methylation percentage/level was calculated as the ratio of the methylated read counts over the sum of both methylated and unmethylated read counts for a single CpG site or across CpGs for a region. The delta methylation was calculated using the log2 transformation of the ratio of methylation level in the B sample (m_B_) and the methylation level in the A sample (m_A_), defined as log_2_ ((m_B_ + 0.1)/(m_A_ + 0.1)) (82). The addition of 0.1 to the numerator and denominator addresses the possibility of zero methylation in one or both samples. Increased methylation in the B sample relative to the A sample is shown as a positive value, whereas decreased methylation in B is shown as a negative value.

### Principal Component Analysis

From the above, we identified 14,037 DMS which we included for further analyses. Principal Component Analysis (PCA) using the prcomp (RRID:SCR_014676) function using R version 4.0.2. ((98) RRID:SCR_001905) was used as a data reduction technique. PCA analyses were carried out for the A samples, the B samples and the changes in methylation (delta methylation values).

### Distance from Transcription Start Site (TSS) and Gene Ontology Analyses were performed using the Genomic Regions Enrichment of Annotations Tool (GREAT)(99)

#### Impact score calculation

Impact scores were calculated using an adapted computational method (100) for calculating polygenic risk scores. From the DMS set, for each individual site, the change in methylation, a binarized QUIC variable (QUIC_bin_) and their interaction were used to predict effortful control at 5 years (EC ∼ “change in methylation by site” * QUIC_bin_). We used a binarized QUIC variable because, although the version used here has 17 items, it was rare for individuals to endorse more than a few. Of the 90 subjects with QUIC and effortful control data, 45% had a raw score of 0, 23% had a raw score of 1, and 32% had a raw score of 2 or more (with values ranging from 2 to 8, reflecting more unpredictable childhoods). The choice to characterize 2 or more as “high” unpredictability provided a sufficient number of individuals to allow for comparison across the two groups.

Sites were then ranked by the p value associated with the interaction coefficient and the top predictive site was added to the set list to be included in the model. The second most predictive site through the last site (using p < 0.05 as a criterion) were then considered sequentially, with each being correlated against sites already included in the set list. Any sites that were not significantly correlated with those already in the list were then added to the set list (Fig. 5A). Genes associated with the selected sites were identified using Genomics Regions Enrichment Annotations Tool (GREAT, RRID:SCR_005807) (99), on the GRCh38 assembly to the single nearest gene within 1000kb. Subject numbers: females = 41, males = 49. 28 ‘low’ females; 13 ‘high’. 16 ‘low’ males; 34 ‘high’.

#### Statistical analyses

All analysis were performed using R 4.0.2 in RStudio (101). Sample preparation and analysis and quality control was performed ‘blind’. Correlations were calculated using Pearson correlation. A comparison of two group means was performed using Student’s t-test. To implement the regression models with interactions, the QUIC scores were converted into binary numbers (QUIC_bin_), with scores greater than one considered ‘high’ and scores of 0 or 1 considered ‘low’ for the reasons described above. Linear regression was performed (EC ∼ QUIC * change in methylation). Figures were made using ggplot2 (RRID:SCR_014601) in R 4.0.2(98). Heatmaps were created in R 4.0.2 using ComplexHeatmap ((102) RRID:SCR_017270) using row normalization. Funding bodies had no role in the study design, data collection or preparation of this manuscript.

## Results

### Methylation profiles distinguish neonatal samples from those obtained at one year of age

To determine how methylation changes may inform future outcomes, two buccal swabs were collected for each infant: the first, during the first month of life and the second at around one year of age (Fig. 1A). In parallel, we collected information on demographics of mothers and infants, including infant sex, birthweight, maternal report of infant ancestry, maternal body mass index pre- and postpartum and income to needs ratio, as well as depressive symptoms. We conducted behavioral assessments of maternal sensitivity as well as measures of unpredictability in the infant’s environment (Table 1). In the infants, we conducted tests of cognitive and emotional development both during the first year of life and at five years of age (Fig. 1A).

**Figure 1.**
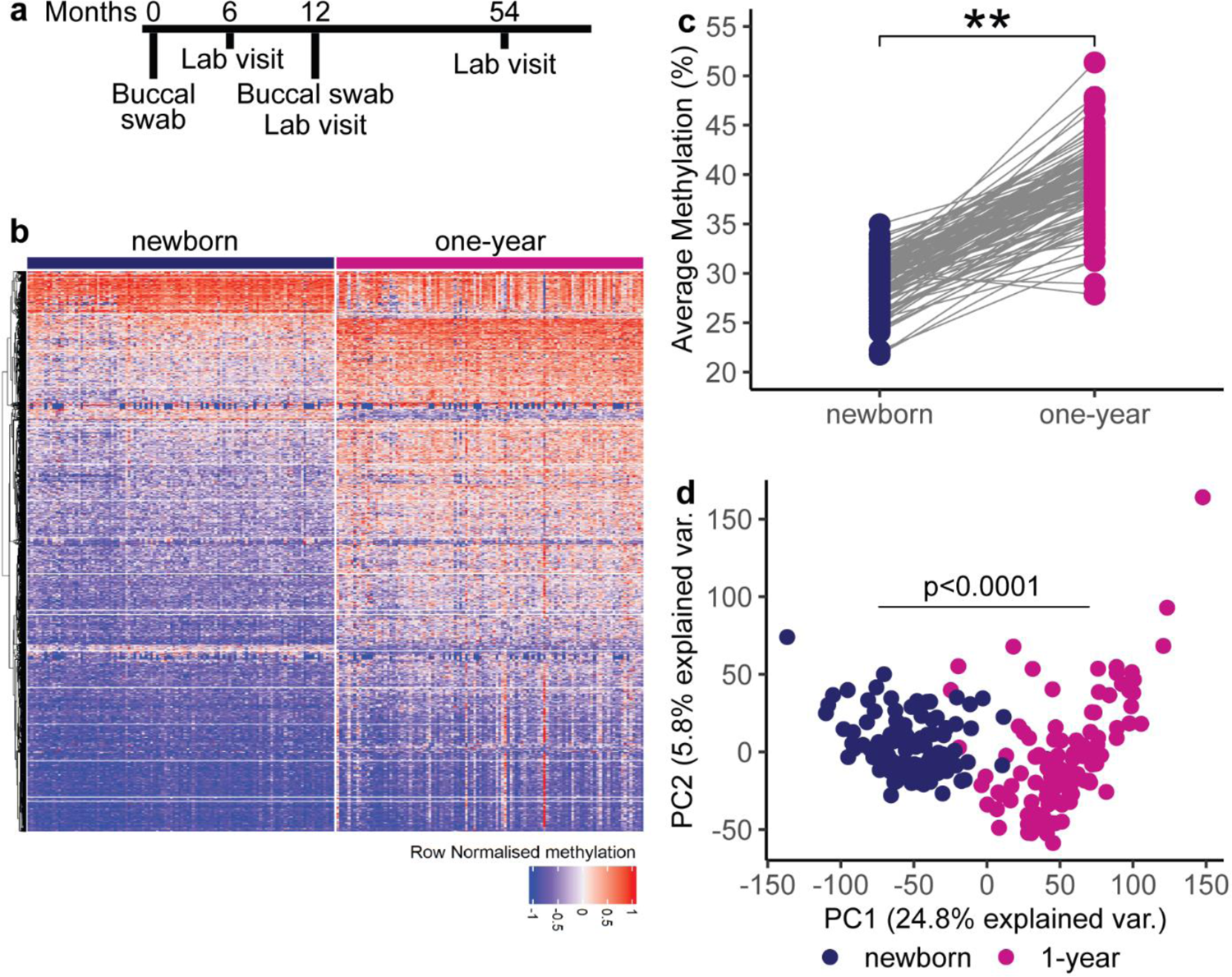
Methylation is influenced by age. A) Timeline of sample collection and assessments in 110 infants. B) Heatmap depicting distinct patterns of methylation distinguishing DNA methylation profiles from newborn and 1-year old children. C) Average percentage methylation at selected sites increases with age. D) Using PCA, the first principal component, explaining 25% of the variance, accounts for the age of sample collection. **p<0.001, bars represent mean, lines represent individual sites.

The analytic workflow of the buccal swab DNA is depicted in the schematic in the Methods section. After mapping each of the two samples from 110 infants to the human genome, we accepted samples with 10 million reads and a coverage of 10 reads or more and identified 1,744,215 methylated sites per newborn sample and 1,743,344 per one-year-old sample that were sequenced with sufficient coverage. We selected for differentially methylated sites by defining them as sites in which methylation was changed by at least ±5% and was significantly different between the newborn and one-year of age samples. We chose a 5% methylation change with the hope that such a change would distinguish the influence of ELA from that of age alone, as found in our preclinical study (82). A site that was identified as having a minimum 5% change in methylation in any individual was then significance-tested for a change in methylation between the A and B samples resulting in a p-value for the i-th site of p_i_. A test was then carried out to determine sites with a significant change from the A to B samples across all individuals by combining the individual test results using −2 ∑^n^_i=1_ ln (*p_i_*) as a test statistic as described in (97). A site that passed a Benjamini-Hochberg false discovery rate of q=0.1 with a cut off = 0.00005 was determined to be a DMS (97). This approach generated 14,037 unique sites that were included for further analysis.

DNA methylation was age dependent (Fig. 1B and 1C), in accord with a robust literature (68,69,75,76,81). There was an average 10% increase in methylation levels for the differentially methylated sites between birth and one year of age (t(_110_)= 31.8, p<0.001), consistent with a previously described general trend for more CpG sites to have increases rather than decreases in DNA methylation during the first year of life (72–75,77). Principal Component Analysis (PCA) of all samples revealed that the first principal component, accounting for 24.8% of the variation in the methylation percentages at the differentially methylated CpG sites, easily distinguished between the samples collected at one month and one year of age (Fig. 1D), in accord with our preclinical study (82) and prior reports (75). Notably, neither sex, nor maternal BMI or infant birthweight separated upon PCA analyses (Supplemental Tables S-2, S-3, S-4 and Fig S-1).

The 14,037 sites differentially methylated between the neonatal and one-year samples resided on all autosomal chromosomes (Fig. 2A). 13,983 of the 14,037 DMS localized to within 1000kb of a transcription start site (TSS; Fig. 2B), and these 13,983 DMS associated with 2,764 unique genes. Gene ontology (GO) analyses demonstrated a striking abundance of genes involved in development (Fig. 2C). We tested the significantly changed sites for overlap with the sites comprising Horvath’s DNA methylation clock as well as the Pediatric epigenetic clock. None of the Horvath clock sites and none of the Pediatric clock sites were among those identified in the current cohort. In contrast, 14 sites corresponding to 8 of the 42 genes identified by Wilkenius et al (75) to distinguish samples from 6 week old from those of 52 week-old Norwegian infants were differentially methylated in our cohort as well. This finding is interesting in view of the homogenous ethnicity and SES status of the 214 Norwegian infants assessed in (75) compared with the diverse ancestry and SES levels in our sample of 110 subjects.

**Figure 2:**
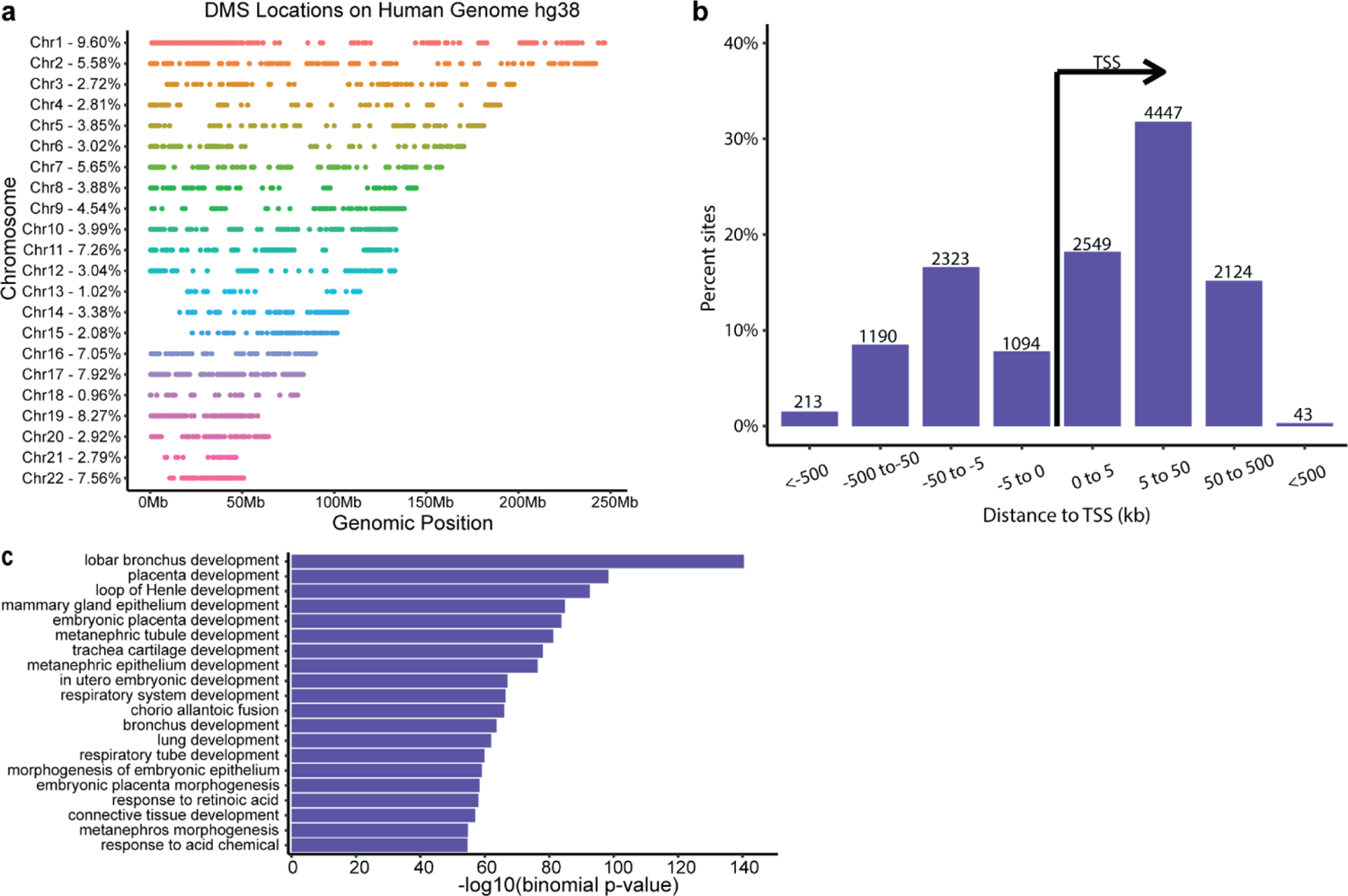
localization and gene ontology of the sites differentially methylated (DMS) between neonatal and one year old samples. (A). Chromosomal distribution of the DMS demonstrates that the reside on all autosomes. Numbers on the left denote the percentage of the overall DMS that localize to each chromosome. (B) Alignment of the DMS with genes and their structures: 13983 of the 14037 DMS localized to within 1000kb of transcription a start site (TSS), and these 13983 DMS associated with 2764 unique genes. (C) Gene ontology identified developmental processes as the key theme of genes associated with DMS between 10-day old and one year old samples of the same child. n=110 infants.

### Methylation changes across time in individual infants, but not methylation at a single time-point, predict effortful control

Calculating the change in methylation at the differentially methylated sites between newborn (first month of life) and one year of age, we performed PCA on the change in methylation (delta methylation) and found that the first component accounted for 8.6% of the variance. Further investigation revealed that the first component correlated highly (R= −0.93, p < 2.2×10^-16^) with the average change in methylation (Fig. 3A). When PCA was performed on the newborn and one-year timepoints separately, the first component also represented the average of the methylation values (R=0.92, p < 1×10^-6^ and R=-0.81, p < 1×10^-6^ respectively). Therefore, further analyses were performed using the average delta methylation values (for analyses involving both time points) and the average methylation values (for analyses involving a single time point).

**Figure 3.**
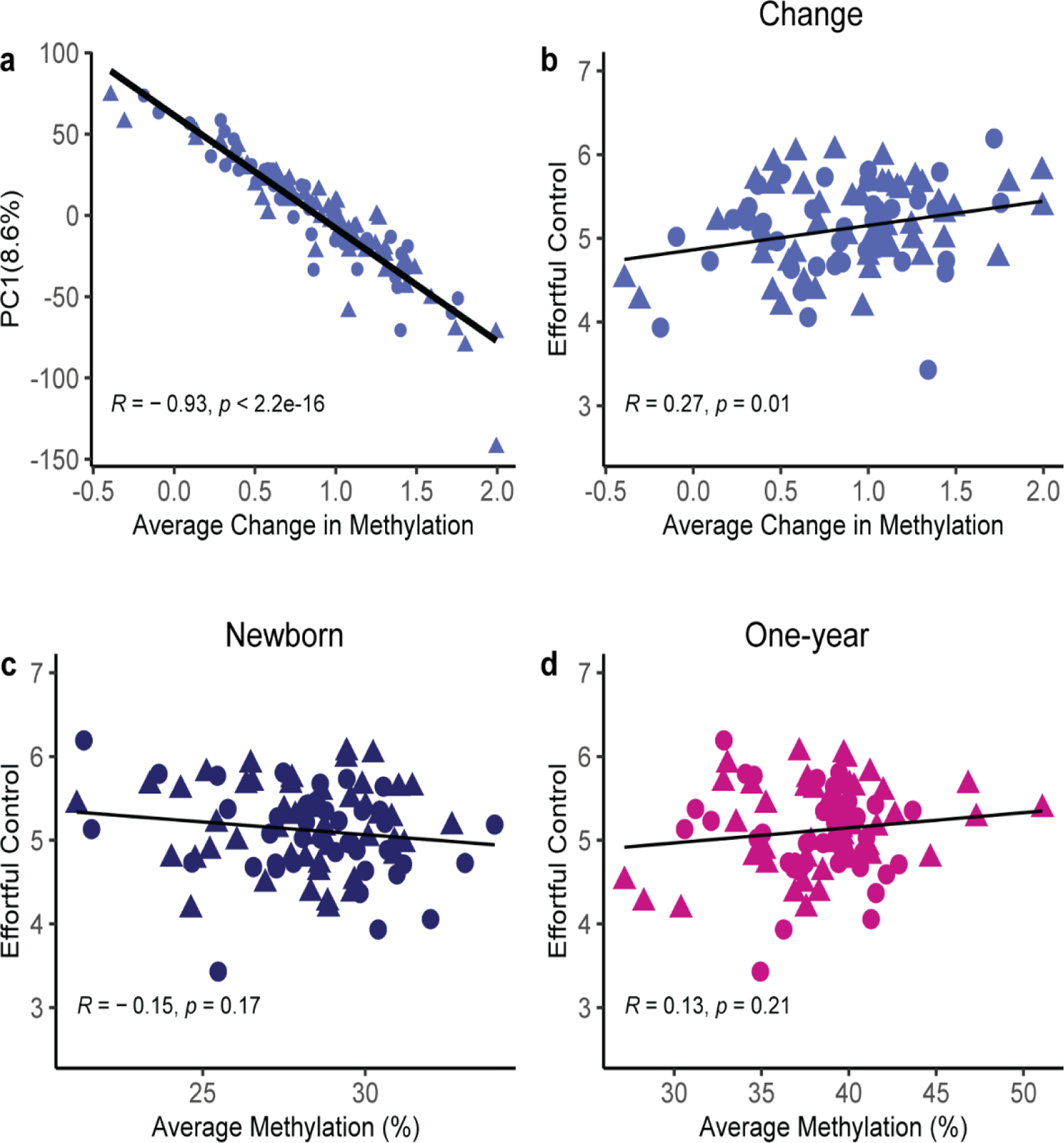
Methylation-changes of individual infants between the ages of 10 days and one year predict effortful control at 5 years. A) The first component of the principal component analysis (PCA) of methylation changes in the ∼ 14,000 differentially methylated sites reflects the average change in methylation (n=110). B) Average methylation change from newborn to one year of age of an individual child predicts effortful control performance at five years of age (n=90). C) Average percent methylation in newborns does not predict outcome. D) Similarly, average percent methylation at one year of age does not predict outcome Note that analogous results were observed when using all 1.74 million methylated sites, as shown in the Supplemental Fig.S-3. Points represent individual samples, circles = females, triangles = males. Line represents linear regression.

We determined the relationship of changes in methylation profiles and effortful control at age five, because effortful control is a reliable predictor of outcomes later in life: lower effortful control in childhood is predictive of poorer mental and physical health, as well as decreased productivity and material success across the lifespan (15,60–63,103–106). The average change in methylation of an individual child from newborn to one year of age was significantly associated with effortful control at 5 years (R= 0.27, p=0.01), accounting for approximately 7.3% of the variance in child effortful control (Fig. 3B). In contrast, analyses of methylation at a single age were not informative: there was no statistically significant association of the average percentage methylation in newborns with effortful control (R=-0.15, p=0.17) (Fig. 3C) and the same applied for the one-year samples (R=0.13, p=0.21) (Fig. 3D). We also computed the relation of average methylation in all 1.7 million methylated CpGs in the neonatal samples and in the one-year samples, and these did not correlate with effortful control (R =-0.069, p=0.52 and R = −0.067, p= 0.53 respectively). In contrast, the average of the changes in methylation of all 1.7 million methylated sites between the one year and neonatal samples correlated significantly with effortful control at age 5 years, though the association was weaker that that observed for the <u>differentially</u> methylated sites (R = 0.22, p=0.036; and see supplemental Figure S-3). Together, these findings suggest that the change in, or “delta” methylation of an individual child over in the first year of life may provide a better indication of the impact of early-life experiences compared with methylation at a single timepoint.

### Unpredictability associates with executive function outcomes at age five years

To determine the dimensions of ELA that might predict effortful control in our cohort, we investigated four established early-life influences on child development: Income-to-needs ratio (INR), maternal depressive symptoms, maternal sensitivity and unpredictability in the child’s caregivers and environment. In our sample, there were weak correlations of INR (R=0.15, p=0.16) (Fig. 4A) and maternal depressive symptoms (R=-0.15, p=0.13) with effortful control (Fig. 4B). There was no correlation of effortful control at age 5 and maternal sensitivity (R=-0.035, p=0.15) (Fig. 3C), and this result obtained also when we analyzed maternal sensitivity separately at 6 and 12 months (R = 0.037. p=0.73 at 6 months and R = −0.086, p=0.45 at 12 months; supplemental Figure S-2). The strongest correlation was observed with unpredictability (R=-0.23, p=0.031) (Fig. 4D), such that high levels of unpredictability associated with lower effortful control, in accord with prior work (10,42). This suggests that unpredictability is a meaningful dimension of ELA and high levels of unpredictability portend poor effortful control at five years of age.

**Figure 4:**
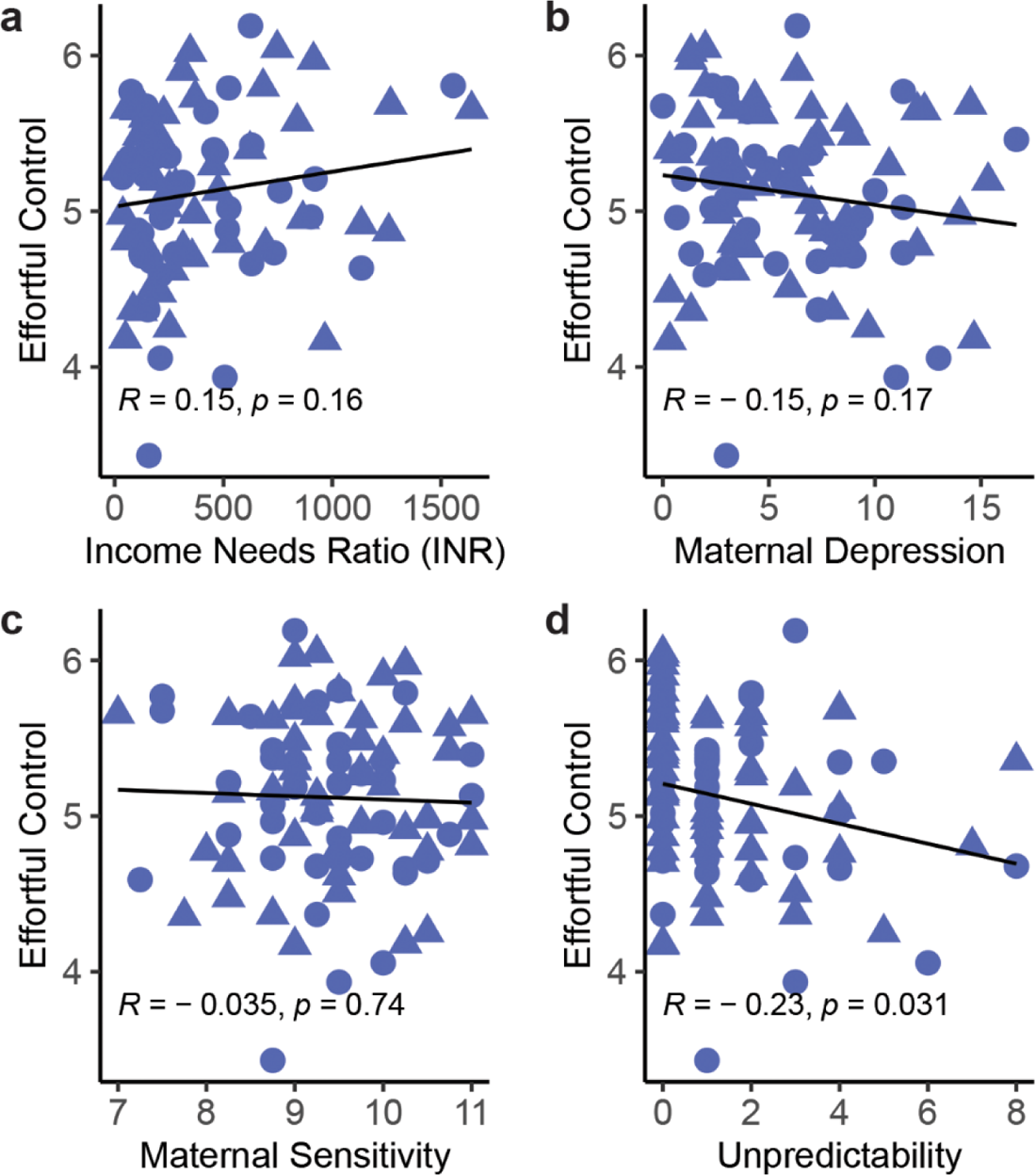
Unpredictability, assessed using the QUIC, portends functional outcomes at 5 years. A) Income / needs ratio (INR) has a weak association with effortful control in our sample of individuals. B) Correlation of maternal depressive symptoms with effort control suggests a weak negative association in these individuals. C) Measures of maternal sensitivity do not correlate with effortful control at 54 months of age in this sample. D) Unpredictability measured using the Questionnaire of Unpredictability in Childhood (QUIC) is inversely correlated with effortful control score at 54 months. Points represent individual samples, circles = females, triangles = males. Line represents linear regression (n=90).

### Unpredictability during the first year of life alters the relation of DNA methylation and later outcomes in girls

Only a subset of individuals experiencing ELA exhibit negative impacts later in life and identifying individuals who are most at risk following ELA is vital to providing targeted interventions. Therefore, we aimed to understand the relationship between maternal and environmental unpredictability, a dimension of ELA, and differential DNA methylation over the first year of life, and probe if these alterations might provide a marker of future deficits in executive function. For the cohort as a whole, we found no direct correlation of the change in methylation over the first year of life with unpredictability (R=-0.07, p=0.51). However, it is well established that sex influences developmental trajectories (107,108), DNA methylation (109,110) and outcomes following ELA, including effortful control (15,71,111,112). Therefore, we analyzed the interaction of DNA methylation and ELA on effortful control, considering sex (Figs. 5A and 5B).

**Figure 5.**
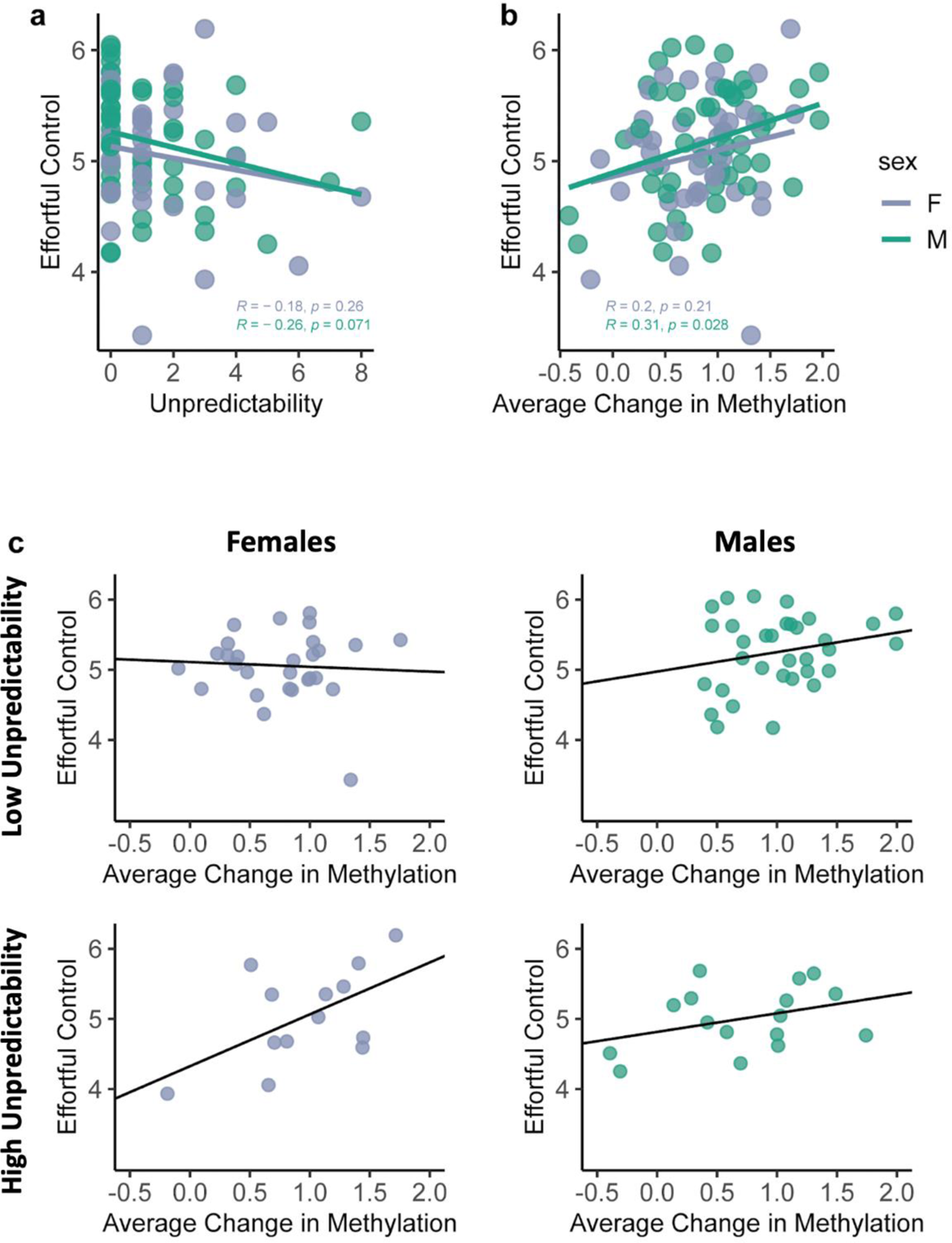
Unpredictability portends child development and may interact with methylation changes over time. A) Unpredictability assessed using the QUIC predicts effortful control at five years of age to a similar degree in both males and females. B) The change in methylation over the first year of life also predicts effortful control at the same age to the same degree in both sexes. n: females=41, males=49. C) There is an interaction between unpredictability and change in methylation in females only: for females who experience high unpredictability, the change in methylation over the first year of life predicts effortful control. In contrast, there is no such interaction observed in males. n: females low=28, females high=13, males low =16, males high=34. Points represent individual samples, purple = females, green = males. Line represents linear regression. F=females, M=Male.

To examine the interaction, individuals were assigned as either experiencing high levels of unpredictability (QUIC >1) or low unpredictability (QUIC ≤1). We used a binary indicator of the QUIC (QUIC_bin_) because of the 90 subjects with QUIC data, two thirds (61) had a raw score of 0 or 1, and only a third (29) had a raw score of 2 or more. Characterizing the latter as experiencing “high” unpredictability provided a sufficient number of individuals to allow for comparison across the two groups. A linear regression model including the indicator of unpredictability, average delta methylation and the interaction, was used to predict effortful control at five years of age for each sex separately and then for both sexes combined (Table 2). In females, there was a significant interaction between the change in methylation and unpredictability (p=0.038), a main effect of unpredictability (p=0.046) and an effect of the average change in methylation (p=0.016) (Fig. 5C). These results suggest that a high level of unpredictability during the first year of life may alter the degree to which changes in differentially methylated sites in females predict effortful control at 5 years. These results were not observed in males: there were no significant interactions or main effects among the parameters tested. Together, these observations imply that sex might influence how unpredictability interacts with changes in DNA methylation and, in girls, this interaction may have a predictive value for risk.

**Table 2:** Regression models testing associations between unpredictability, methylation and effortful control at 5 years of age.

| Model | Predictors | Estimates | CI | p |
| --- | --- | --- | --- | --- |
| *Combined sexes* |  |  |  |  |
|  | (intercept) | 4.672 | 4.504-4.840 | <0.0001 |
|  | mean change in methylation | 0.423 | 0.256-0.590 | 0.013 |
|  | QUIC[low] | 0.337 | 0.113-0.562 | 0.136 |
|  | Interaction | -0.250 | -0.474- -0.027 | 0.266 |
| *Girls* |  |  |  |  |
|  | (intercept) | 4.327 | 3.681-4.973 | <0.0001 |
|  | mean change in methylation | 0.740 | 0.147-1.333 | 0.016 |
|  | QUIC[low] | 0.784 | 0.015-1.553 | 0.046 |
|  | Interaction | -0.810 | -1.570—0.048 | 0.038 |
| *Boys* |  |  |  |  |
|  | (intercept) | 4.815 | 4.621-5.001 | <0.0001 |
|  | mean change in methylation | 0.266 | 0.059-0.473 | 0.206 |
|  | QUIC[low] | 0.161 | -0.134-0.459 | 0.588 |
|  | Interaction | 0.011 | -0.279-0.301 | 0.971 |

### Calculating an impact score as a predictive marker of individuals susceptible to early-life adversity

The association of average change in DNA methylation and unpredictability with executive function at age five led us to consider an approach for identifying the potential contributions of specific sites (and their respective genes) to the overall relationship. Combined, such strong-effect sites might serve to construct predictive (“polyepigenic”) risk scores for vulnerability to adverse outcomes. Whereas risk scores typically require cohorts with a minimum of 100 subjects (113), these size recommendations are based on the use of each subject as a unitary entity within a population. However, here, the ‘delta methylation’ approach compares each individual to themselves, reducing the effect of population variance on diluting effect sizes.

We used the data from females because the interactions were present in female samples only. Adopting a polygenic risk score clumping and thresholding method (100,113), we ran a linear regression using each of the differentially methylated sites and the binarized QUIC score to predict effortful control, and assessed the significance of the interaction (Fig. 6A). The algorithm identified 37 ‘significant’ sites (p<0.05) (Fig. 6B and Table 3). By summing the change in methylation from birth to one year of age at these 37 sites in females, we created an impact score. We analyzed the females who had experience greater and lesser degrees of unpredictability separately. The impact score significantly predicted an individual girl’s effortful control performance at 5 years of age (score x QUIC_bin_ interaction; R^2^= 0.20, p=0.0016) (Fig. 6C) in females who had experienced greater unpredictability. Note that the significance of the interaction here is expected given the way in which specific sites were selected. In this cohort of females, this algorithm predicted individuals who would develop poor effortful control in later years.

**Figure 6.**
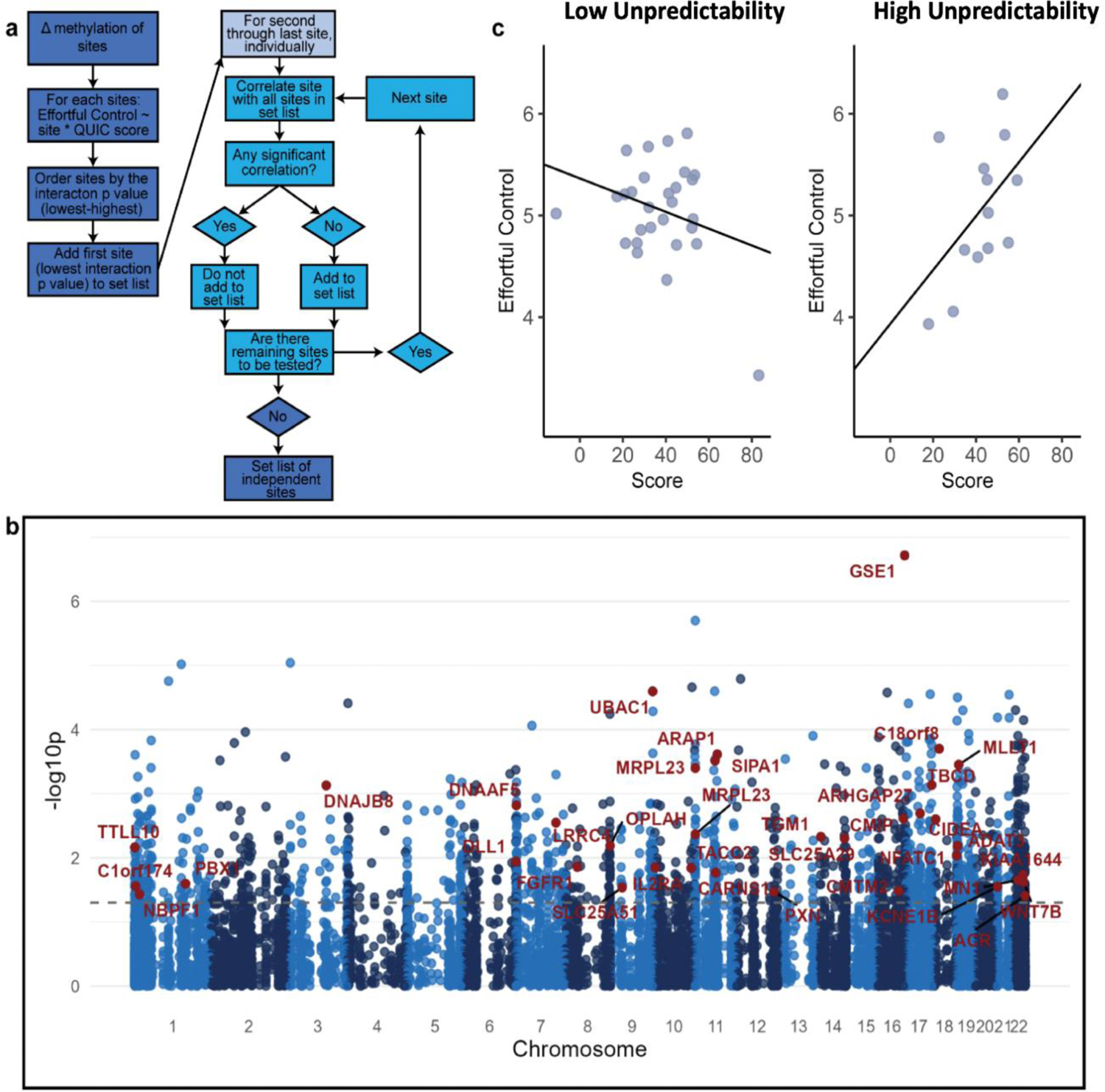
Impact scores identify individuals vulnerable to poor outcomes following the unpredictability dimension of early-life adversity. A) Flow chart of the computation method used to select highly contributing sites. The clumping and thresholding method identifies sites that appear to have a significant interaction but do not correlate highly with other sites that have already been selected. B) Manhattan plot representing the differentially methylated sites (blue) and the distribution across the chromosomes and the corresponding significance score (-log10p) of each site interacting with QUIC to predict effortful control in females. Sites in red are those selected via the clumping and thresholding algorithm. Dotted line is at p=0.05. C) The significant interaction of QUIC and risk score calculated from top sites according to our model predicts effortful control at 5 years in females who have experienced more adversity n: low=28, high=13. Line represents linear regression.

**Table 3.** Top sites contributing to the impact score.

| Gene | Site | p value | R<sup>2</sup> |
| --- | --- | --- | --- |
| GSE1 | chr16:85650857-85650858 | 1.92E-07 | 0.49 |
| UBAC1 | chr9:136017659-136017660 | 2.54E-05 | 0.34 |
| C18orf8 | chr18:23524804-23524805 | 1.99E-04 | 0.27 |
| ARAP1 | chr11:72754843-72754844 | 2.43E-04 | 0.27 |
| SIPA1 | chr11:65642488-65642489 | 3.05E-04 | 0.25 |
| MLLT1 | chr19:6270257-6270258 | 3.53E-04 | 0.28 |
| MRPL23 | chr11:1951274-1951275 | 3.97E-04 | 0.24 |
| TBCD | chr17:82795360-82795361 | 7.34E-04 | 0.22 |
| DNAJB8 | chr3:128421485-128421486 | 7.43E-04 | 0.25 |
| DNAAF5 | chr7:779363-779364 | 1.52E-03 | 0.18 |
| ARHGAP27 | chr17:45405991-45405992 | 2.03E-03 | 0.18 |
| CMIP | chr16:81431947-81431948 | 2.44E-03 | 0.17 |
| CIDEA | chr18:12278873-12278874 | 2.51E-03 | 0.16 |
| LRRC4 | chr7:128030495-128030496 | 2.84E-03 | 0.16 |
| MRPL23 | chr11:1951752-1951753 | 4.30E-03 | 0.20 |
| TGM1 | chr14:24259744-24259745 | 4.70E-03 | 0.13 |
| SLC25A29 | chr14:100297661-100297662 | 4.97E-03 | 0.13 |
| OPLAH | chr8:144052079-144052080 | 6.48E-03 | 0.12 |
| ADAT3 | chr19:1912103-1912104 | 6.50E-03 | 0.12 |
| TTLL10 | chr1:1145182-1145183 | 6.88E-03 | 0.15 |
| NFATC1 | chr18:79485573-79485574 | 9.21E-03 | 0.12 |
| DLL1 | chr6:170227779-170227780 | 1.15E-02 | 0.11 |
| FGFR1 | chr8:38554659-38554660 | 1.38E-02 | 0.10 |
| IL2RA | chr10:6036875-6036876 | 1.41E-02 | 0.09 |
| TACC2 | chr10:122085904-122085905 | 1.44E-02 | 0.11 |
| CARNS1 | chr11:67411863-67411864 | 1.71E-02 | 0.14 |
| KIAA1644 | chr22:44286585-44286586 | 1.84E-02 | 0.07 |
| MN1 | chr22:27797185-27797186 | 2.22E-02 | 0.14 |
| WNT7B | chr22:45972341-45972342 | 2.38E-02 | 0.10 |
| PBX1 | chr1:164576643-164576644 | 2.53E-02 | 0.12 |
| C1orf174 | chr1:4161521-4161522 | 2.78E-02 | 0.10 |
| KCNE1B | chr21:8434920-8434921 | 2.83E-02 | 0.10 |
| SLC25A51 | chr9:37938671-37938672 | 2.90E-02 | 0.08 |
| CMTM2 | chr16:66579316-66579317 | 3.28E-02 | 0.05 |
| PXN | chr12:120263326-120263327 | 3.39E-02 | 0.09 |
| NBPF1 | chr1:16725298-16725299 | 3.73E-02 | 0.18 |
| ACR | chr22:50730818-50730819 | 3.90E-02 | 0.09 |

The 37 DMS comprising the impact score belonged to 36 genes. GO analyses and gene network analyses revealed no significantly enriched terms. In addition, comparing these sites to sites implicated in the Horvath (69) and Pediatric epigenetic clocks (76) uncovered no overlapping sites. Thus, in a small female cohort, we identified an approach that may have the potential, in future large validation studies, to provide a predictive marker of individuals at risk of worse outcomes following high levels of unpredictability and, possibly, other dimensions of ELA.

## Discussion

The principal findings presented here are: 1) The use of a longitudinal, within-subject approach identifies changes in methylation over the first postnatal year as a feasible tool which may allow better prediction of executive function at age five years compared to methylation profiles at a single time point. 2) Exposures to a higher degree of unpredictability in early life, a dimension of ELA, correlate with poorer effortful control. 3) The interaction of this dimension of ELA with changes in methylation is sex-dependent in our cohort. 4) In girls, unpredictability interacts with change in methylation to presage effortful control, suggesting that unpredictability in early life may alter the relationship of DNA methylation and outcomes later in childhood. 5) A tentative impact score was created using the change in methylation in girls, aiming to provide a predictive marker of the influence of high levels of early-life unpredictability on the future outcome of an individual child. This score should be validated in future studies as a potential indicator of risk.

Identifying individuals with a high risk of developing cognitive and emotional problems after sustaining pre-or early postnatal adversity is important: such a discovery will allow targeting preventative and interventional strategies to those who need them most. Indeed, a number of investigative groups and consortia have aimed to employ DNA methylation profiles of blood or buccal swab cells of infants and children as a correlate of ELA and a predictor of the subsequent outcomes (73,74,76,77,79,114–116).

DNA methylation levels vary with age, and normative patterns and rates of these changes have been established, providing epigenetic--or DNA methylation (DNAm)—clocks (67,76). Deviations from this ‘clock’, and especially acceleration of DNAm vs chronological ages, have been considered predictive of aging and disease (68). The rate of DNA methylation change is especially rapid early in life, and a body of work has focused on creating and harmonizing DNAm clocks that are optimal for infants and children (71,72,75–77,79), and on the influence of ELA on modulating and especially accelerating the pediatric epigenetic clocks (73,74,76,79). Additional clocks have been identified for specific early-life epochs including the use of cord blood to assess gestational age (72) and buccal swabs to probe the first year of life (75). More recently, Fang et al., 2023 compared seven pediatric clocks, highlighting the heterogeneity of sites identified across studies: Indeed, of 2,587 CpGs, 2,206 (>80%) were specific to only one clock (117). Thus, the modest overlap of sites identified here with those identified in other cohorts is not surprising.

We find that the changes in overall average methylation between one-year old and neonatal samples for sites with a minimum 5% change is largely positive (higher methylation). The methylation changes that happen over time during childhood are bi-directional (75,76,81). Focusing on the neonatal period, Wilkenius et al., found increased methylation in 36 of 42 sites that change significantly during the first postnatal year, in accord with the current study (75). They considered this augmented methylation surprising because higher methylation tends to predict reduced gene expression. These authors speculated that the putative reduction in gene expression during the first year of life might be compensatory to explosive gene expression *in utero* (75). It is unclear whether or not the overall increase in methylation observed in the current study is beneficial. While it might be considered an acceleration of the epigenetic ‘clock’ for this age, the implications of such acceleration are not obvious: Whereas in adults, acceleration of epigenomic clocks is almost universally detrimental and associated with accelerated ageing, disease and death, Suarez er al., (73) identified a <u>decelerated</u> DNA-methylation clock in boys with mental health problems following prenatal exposure to maternal anxiety. Indeed, some authors suggest that DNA-methylation clock acceleration in childhood implies accelerated physical and mental development, with salubrious consequences. Clearly, additional empiric information and longitudinal studies are required to address these intriguing questions.

The majority of studies to date have employed profiles of DNA methylation in samples obtained at a single time point. Dunn et al., (118) and Lussier et al. (119) used longitudinal approaches, focusing specifically on the timing of and dimension of ELA in influencing methylation profiles. The group identified ages 3-5 as putative sensitive period for methylation effects of ELA. However, these studies did not assess ELA during the first postnatal year. The current longitudinal study suggests that the first year of life may also be an important sensitive period for the effects of some dimensions of ELA (e.g., unpredictability) on DNA methylation as well as on neurodevelopmental outcomes.

Specifically, we aimed here to minimize large variances in methylation among individuals, account for the change of methylation with age (73,74,82,118,120–122), and avoid dilution of the potential effects of neonatal ELA over a lifetime. Therefore. we employed a longitudinal, or ‘within subject’ approach (82,120,123–125) and sampled infants at a relatively short interval--one year--which mitigated the potential dilution of an epigenomic ‘signature’ of ELA by subsequent life events. Finally, we examined for the well-established effects of age on methylation profiles, employing both Horvath’s DNA methylation (DNAm) clock as well as the more recently described Pediatric epigenomic clock (76). We found that the change in methylation profile of an individual child between the first month of life and one year of age was superior at predicting neurodevelopment at five years compared with a methylation profile derived from a single timepoint.

In the current analysis of a cohort of 110 children, we quantified several types of adversity during the interval year between the two samples and examined the relative contribution of these measures of ELA to both changes in DNA methylation during the first year of life and to effortful control at age five years. In addition to poverty (assessed as income-to-needs ratio), and maternal depressive symptoms and sensitivity, we tested the role of unpredictable signals from the mother/caretaker and the household environment. This dimension of adversity has emerged in our work (37,39–41,43–45,47,126,127) and independently, in work by others (42,128–130) as a contributor to cognitive outcomes including effortful control (39–41,131) and emotional (41,44,127,130,132) outcomes in children, adolescents and adults. The neurobiological basis for the detrimental effects of unpredictable environmental signals on brain development are not fully understood. In both humans and experimental models, sensory input from the environment (e.g., light patterns, patterns of tones) are required for appropriate maturation of the respective brain circuits. In experimental models, unpredictable patterns of sensory inputs may impact brain circuit maturation by disrupting the selective microglial pruning of synapses (30,133).

In the relatively small cohort assessed here, both lesser changes in methylation during the first year of life and high levels of maternal unpredictability were predictive of poorer effortful control at age five years. In addition, further analyses suggested sex-dependent differences. Interactions of methylation changes and unpredictability were observed, but only in girls. The discovery of sex effects of ELA on methylation profiles and outcome already prior to puberty is intriguing. In adults, but not for children, a more rapid epigenetic ageing has been reported in women (134,135), whereas others found a more rapid epigenetic ageing in adolescent girls with a history of ELA, but not in boys (136). While many sex-differences in post-pubertal individuals are ascribed to sex hormones, the effects of sex in both the current study and others (71,73,79), suggest other biological differences between males and females that are at play already soon after birth.

The basis of these sex differences remains enigmatic: Dammering et al., who studied the effects of postnatal ELA on DNA methylation in the context of ‘Epigenetic Ageing’ found a greater epigenetic ageing in girls than in boys (79). In contrast, looking at the effect of prenatal exposure to maternal depression on DNA-methylation age of newborns, Suarez et al., identified an effect in boys but not girls (73), and a similar male vulnerability was observed by McGill et al., for maternal anxiety (71), and by work from our group (140). Studies of selective vulnerabilities of males to prenatal stress are buttressed by work in experimental models demonstrating similar male vulnerability (137). For postnatal stress, both Dammering et al. (79) and our own studies (59) suggest greater vulnerability in girls. Together, the combined body of work suggests that sex effects can be detected prior to puberty, and that different types of adversity and its timing, i.e., the developmental age in which adversity takes place, may influence which sex is more affected.

Comparing the current work to other studies, we note our use of reduced representation bisulfite sequencing (RRBS). Other groups (121,122,124,125) have employed bisulfite conversion and genomic DNA methylation profiling using the Illumina HumanMethylation450 BeadChip which assesses DNA methylation levels at >480,000 CpG sites. Still others used targeted sequencing of specific sites (123). All these methods have assets and limitations: RRBS only samples ∼5% of the genome, but includes ∼95% of gene-related CpG sites. In our hands, it uncovered methylation at ∼1.74 million CpGs, well in the range of the Illumina chip approach. In contrast, the use of methylation panels allows sequencing assessment of predefined sites, yet does not allow discovery of novel sites as markers. Hence, we believe RRBS provides a compromise between targeted sequencing and a whole genome approach.

The discovery of a robust association of unpredictability, changes in methylation and effortful control in girls led us to probe whether specific differentially methylated sites could be used to create an individually predictive impact score. While general recommendations in the literature suggest against generating genetic or epigenetic risk scores to cohorts smaller than 100 samples (113), these size recommendations are based on the use of each subject as a unitary entity within a population. However, here, the ‘delta methylation’ approach compares each individual to themselves, reducing the effect of population variance on diluting effect sizes, perhaps analogous to the use evoked potentials compared with EEG (138,139).

There are several limitations to the current study, the primary being the cohort size. Epigenetic and genetic studies often include tens or hundreds of thousands of subjects, providing power that our cohort of 110 infants does not permit. In addition, parsing the group by sex further reduces sample size, with a risk of overfitting. We acknowledge this issue and note that studies of similar size can provide important and innovative information. For example Jovanovic et al. (74) uncovered an important effect of ELA on DNA methylation in a cohort of 101 subjects.

In addition, we aim here to address the cohort size caveat in part by the use of a longitudinal within-subject design, enabling assessment of DNA changes within an individual over time rather than a cross section comparison of different groups, which is more sensitive to random effects and overfitting in small samples. Capitalizing on the within subject design, we attempt to generate a polyepigenetic impact score, and note that this score has yet to be validated because the size of the current cohort did not permit splitting it into training and testing subsets (113). Thus, validation of the current impact score requires larger naïve datasets. Nevertherless, we suggest that the technologies and approaches presented here provide valuable insights into the potential of using the interaction of early-life adversity and methylation changes across defined epochs as potential indicators of the impact (‘epigenetic scar’) of adversity on an individual child, with significant predictive promise.

## Data Sharing Statement

Data sharing will comply with National Institute of Health guidelines. Epigenomic datasets will be submitted to the appropriate databanks and will be made available on request.

## Acknowledgments

The authors thank the families who participated in this project and our dedicated staff.

## Contributors

AKS: Conceptualization, data curation, formal analysis, investigation, methodology, software, validation, visualization, writing - original draft

RW: Data curation, formal analysis, investigation, methodology, software, validation, visualization

NK: Data curation, formal analysis, investigation, methodology

CWT: Conceptualization, data curation, formal analysis, investigation, methodology, software, validation

HA: Software, validation, methodology

AM: Conceptualization, funding acquisition, methodology, validation, project administration, resources, supervision, writing - review and editing

HSS: Conceptualization, funding acquisition, methodology, validation, project administration, resources, supervision, writing - review and editing

LG: Conceptualization, funding acquisition, methodology, validation, project administration, resources, supervision, writing - review and editing

TZB: Conceptualization, funding acquisition, methodology, validation, project administration, resources, supervision, writing, revisions, review and editing

