## supplemental material for "Within-subject changes in methylome profile identify individual signatures of early-life adversity, with a potential to predict neuropsychiatric outcome"

**Supplemental information:**

**Supplemental Table S-1:**

**The Questionnaire on Unpredictability Iin Childhood (QUIC): Questions used in the current study and endorsement rates for each**

| Question | Percent Endorsement |
|---|---|
| At least one of my child's parents kept track of what my child ate (e.g., made sure that he/she did not skip meals or tried to make sure he/she ate healthy food). (**R**) | 100 |
| My child's family ate a meal together most days. (**R**) | 95.8 |
| At least one of my child's parents tried to make sure he/she got a good night's sleep (e.g., he/she had a regular bedtime, parents checked to make sure he/she went to sleep). (**R**) | 100 |
| My child had a bedtime routine (e.g., was tucked in, parents read him/her a book, took a bath). (**R**) | 94.8 |
| One or more of my child's parents had punishments that were unpredictable. | 15.6 |
| My child often wondered whether or not one of his/her parents would come home at the end of the day. | 7.3 |
| There were often people coming and going in my child's house that he/she did not expect to be there. | 3.1 |
| At least one of my child's parents made time each day to see how my child was doing. (**R**) | 100 |
| My child's parents had a stable relationship with each other. (**R**) | 22.7 |
| One or more of my child's parents was disorganized | 22.9 |
| One or more of my child's parents was unpredictable. | 11.5 |
| When one or more of my child's parents was upset, my child did not know how they would act. | 5.2 |
| One or more of my child's parents could go from calm to furious in an instant. | 18.8 |
| One or more of my child's parents could go from calm to stressed or nervous in an instant. | 33.3 |
| My child lived in a clean house. (**R**) | 97.9 |
| My child lived in a cluttered house (e.g., piles of stuff everywhere). | 8.3 |
| In my child's house, things he/she needed were often misplaced so that we could not find them. | 3.1 |
| Note: R indicates a reverse-scored item | |

**Supplemental Table S-2**:

**Analysis of infant birth weight correlation with principal components obtained from the differentially methylated sites (DMS) identified in the study.**

| PC | Percent_variance_explained | R | p_value |
|---|---|---|---|
| PC1 | 8.61 | 0.162 | 0.091 |
| PC2 | 4.57 | 0.027 | 0.777 |
| PC3 | 2.89 | -0.129 | 0.18 |
| PC4 | 2.64 | 0.042 | 0.664 |
| PC5 | 2.36 | 0.03 | 0.755 |
| PC6 | 2.18 | 0.042 | 0.666 |
| PC7 | 1.91 | -0.075 | 0.437 |
| PC8 | 1.84 | 0.014 | 0.885 |
| PC9 | 1.81 | 0.068 | 0.48 |
| PC10 | 1.64 | -0.014 | 0.888 |
| PC11 | 1.59 | 0.097 | 0.313 |
| PC12 | 1.59 | 0.039 | 0.687 |
| PC13 | 1.54 | 0.081 | 0.401 |
| PC14 | 1.52 | 0.052 | 0.59 |
| PC15 | 1.44 | 0.119 | 0.214 |
| PC16 | 1.36 | -0.01 | 0.92 |
| PC17 | 1.31 | 0.051 | 0.593 |
| PC18 | 1.28 | 0.031 | 0.748 |
| PC19 | 1.22 | -0.078 | 0.415 |
| PC20 | 1.19 | 0.254 | 0.007 |

**Supplemental Table S-3:**

**Analysis of Maternal (postpartum) body mass index (BMI) correlation with principal components obtained from the differentially methylated sites (DMS) identified in the study**

| PC | %_of_variance_explained | R | pvalue |
|---|---|---|---|
| PC1 | 8.61 | -0.152 | 0.164 |
| PC2 | 4.57 | 0.009 | 0.936 |
| PC3 | 2.89 | -0.068 | 0.536 |
| PC4 | 2.64 | -0.005 | 0.962 |
| PC5 | 2.36 | 0.154 | 0.158 |
| PC6 | 2.18 | -0.02 | 0.858 |
| PC7 | 1.91 | 0.098 | 0.369 |
| PC8 | 1.84 | -0.072 | 0.51 |
| PC9 | 1.81 | 0.174 | 0.108 |
| PC10 | 1.64 | 0.066 | 0.545 |
| PC11 | 1.59 | 0.004 | 0.973 |
| PC12 | 1.59 | -0.009 | 0.936 |
| PC13 | 1.54 | -0.067 | 0.541 |
| PC14 | 1.52 | -0.052 | 0.634 |
| PC15 | 1.44 | -0.073 | 0.504 |
| PC16 | 1.36 | -0.084 | 0.441 |
| PC17 | 1.31 | 0.052 | 0.633 |
| PC18 | 1.28 | -0.247 | 0.022 |
| PC19 | 1.22 | 0.04 | 0.717 |
| PC20 | 1.19 | 0.026 | 0.811 |

**Supplemental Table S-4**:

**Analysis of Maternal (pre-partum) body mass index (BMI) correlation with principal components obtained from the differentially methylated sites (DMS) identified in the study**

| PC | %_of_variance_explained | R | pvalue |
|---|---|---|---|
| PC1 | 8.61 | -0.195 | 0.056 |
| PC2 | 4.57 | 0.089 | 0.385 |
| PC3 | 2.89 | -0.083 | 0.419 |
| PC4 | 2.64 | -0.051 | 0.623 |
| PC5 | 2.36 | 0.133 | 0.193 |
| PC6 | 2.18 | -0.113 | 0.272 |
| PC7 | 1.91 | 0.057 | 0.583 |
| PC8 | 1.84 | -0.085 | 0.406 |
| PC9 | 1.81 | 0.133 | 0.193 |
| PC10 | 1.64 | 0.131 | 0.201 |
| PC11 | 1.59 | -0.08 | 0.437 |
| PC12 | 1.59 | -0.033 | 0.75 |
| PC13 | 1.54 | -0.048 | 0.639 |
| PC14 | 1.52 | -0.026 | 0.797 |
| PC15 | 1.44 | -0.042 | 0.685 |
| PC16 | 1.36 | -0.147 | 0.15 |
| PC17 | 1.31 | 0.057 | 0.583 |
| PC18 | 1.28 | -0.107 | 0.296 |
| PC19 | 1.22 | 0.255 | 0.012 |
| PC20 | 1.19 | -0.154 | 0.133 |

**Supplemental Figure S-1**

**Scatterplots showing first two principal components derived from methylated sites and infant sex for a single time-point (left) and first two principal components derived from the sites differentially methylated between the neonatal and one year samples (DMS) and infant sex (right).**

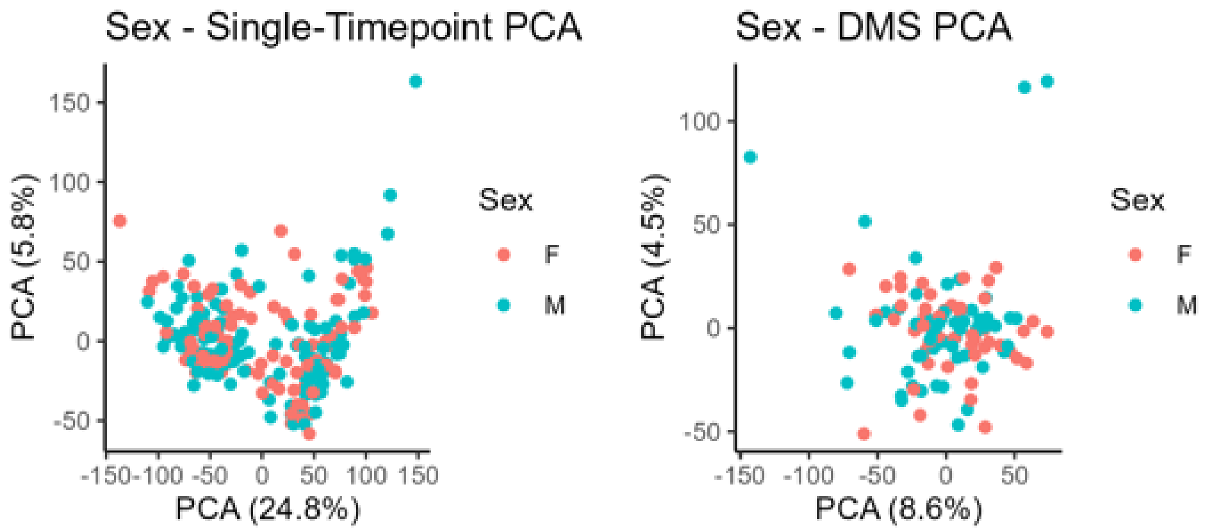


Legend: The procedures for calling of methylated sites and those differentially methylated between the neonatal and one-year old sample of an individual child are described in the Methods section. As shown in the graphs, sex did not segregate with the first two principal components of either methylated sites identified at a single time point or the ~14,000 sites that were differentially methylated between the two time-points. N = 110

**Supplemental Figure S-2**

**Analysis of the correlation of maternal sensitivity at 6 months and 12 months with effortful control of her child at age 5 years.**

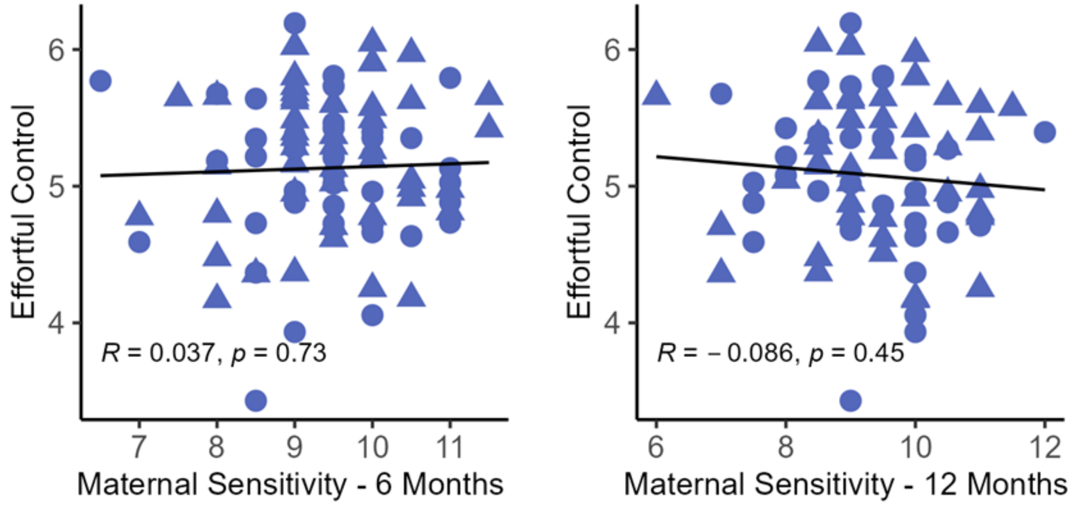

Legend: The procedures for assessing maternal sensitivity are described in the Methods section. As shown in the graphs, maternal sensitivity at infant age of 6 months (left) or 12 months (right) did not correlate with the child's effortful control at age five years. The correlation of the averaged maternal sensitivity with effortful control is shown in Main Figure 3. N = 90.

**Supplemental Figure S-3**

**The correlation of the average methylation of all methylated sites at one month (left), or at age one year (center), as well as the average change in methylation for all methylated sites between the neonatal and one-year samples of an individual child (right), with effortful control at age 54 months.**

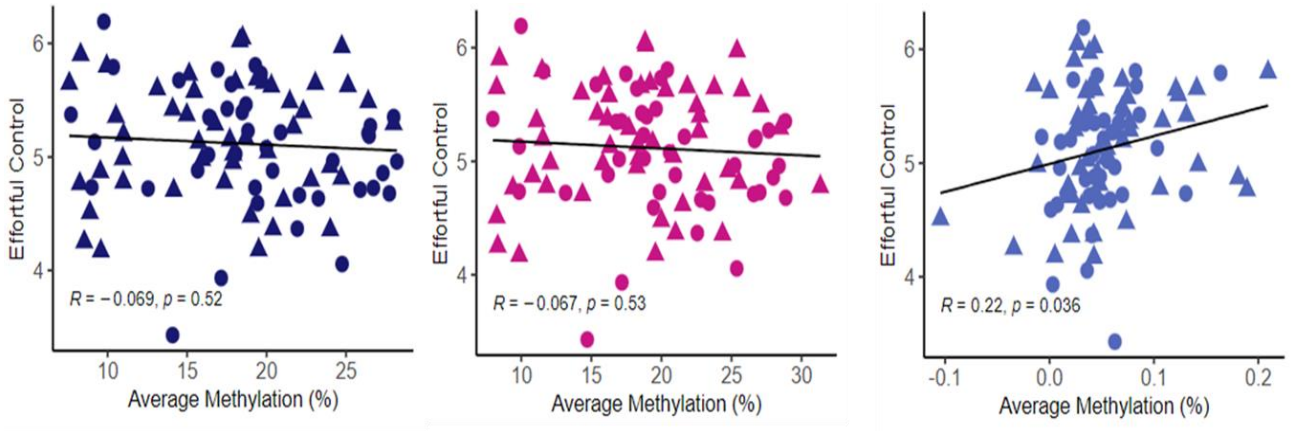


Legend: The procedures for identifying methylated sites and differentially methylated sites (DMS) are described in the Methods section. As shown in the graphs, the correlation of all ~1.74 million methylated sites identified at a single time point (one month on the left or one year in the center panel) did not correlate with future effortful control. In contrast, average change in methylation between the neonatal and one-year samples of the ~1.74 million methylated sites for an individual child correlated highly with effortful control at age 54 months. N = 90.
